# An auxin-inducible degron system for trypanosomes

**DOI:** 10.1101/2025.06.17.660065

**Authors:** Alexandra Klein, Alice McDowell, Christian R S Reis, Rodrigo Pontes de Lima, Danielle M N Moura, Janaina de Freitas Nascimento, Osvaldo P de Melo Neto, Mark Carrington

## Abstract

The ability to specifically degrade one protein in a cell provides an immediate insight into any resulting phenotype and thus function that may be occluded by secondary effects using other techniques. The auxin inducible degron is a method developed to achieve this end and is effective in mammalian and yeast cells. The approach is dependent on the recognition of a tagged target protein by an endogenous E3 ubiquitin ligase modified by the expression of a plant F-box protein. This approach was initially unsuccessful in trypanosomes probably due to a lack of interaction between the endogenous E3 ubiquitin ligase and the plant F-box protein. This was overcome by expressing the entire rice E3 ubiquitin ligase in trypanosomes and here we describe the production of the auxin inducible degron competent cell lines and tagging of target genes.

## 1 Introduction

The ability to specifically ablate a protein still underlies the majority of investigations into the function of a particular gene. The first approach used to identify a role for a gene in a process involved making null or conditional mutants through random mutagenesis and screening for a particular phenotype. This type of forward genetics, in various guises, remains a powerful tool as it requires no knowledge of the identity of the gene. The advent of widely available genome sequences allowed the identification of target genes that may or may not be involved in a process of interest and led to an explosion in reverse genetics. In this approach, the gene was identified from sequence, altered in some way, and the resulting phenotype investigated. Any phenotype usually results from the loss of a protein in a cell and this can be achieved at three levels: deletion of the gene, ablation of the mRNA by RNAi, or targeted degradation of the protein. All are effective but each has disadvantages; recovery after gene deletion may result in a compensatory change that masks a phenotype. RNAi based degradation of a specific mRNA occurs rapidly but not always completely, also the protein lingers on and often has to drop below a threshold before a phenotype becomes apparent. This makes it difficult to separate primary and secondary phenotypes. Protein ablation through targeted degradation is not always complete but is rapid and provides an opportunity to identify a primary phenotype.

The African trypanosome *Trypanosoma brucei* is the causal agent of disease in humans, livestock and wild animals. It has been intensively studied for more than 100 years and research now centres on both the causes of pathology and on basic cellular and molecular processes of a divergent eukaryote. Forward genetics is non-trivial as trypanosomes are diploid, crosses occur in the salivary glands of the tsetse fly and are a non-obligatory part of the life cycle. The genome was sequenced 20 years ago [1] and this opened up the use of reverse genetics as an approach to address questions about trypanosome biology. The dominant approach has been RNAi [2] which has been successful both in identifying the role of individual genes and has been modified for forward genetics using whole genome RNAi libraries [3,4]. The RNAi approach usually uses tetracycline-inducible promoters to produce double stranded RNA and the target mRNA is dramatically reduced within one hour [2]. However, in many experiments a phenotype does not appear for another 24 hours or more (for example [5]) and this has been interpreted as the time taken for the protein of interest to drop below a threshold by dilution resulting from cell proliferation. This lag means it can be difficult to separate primary and secondary phenotypes.

The time taken for protein levels to diminish is a particular problem in experiments investigating mRNA turnover as the half-life of mRNAs in trypanosomes ranges from a few minutes to a couple of hours [6]. The time taken for protein depletion after RNAi induction means it is difficult to resolve which mRNAs are altered as a primary effect of the RNAi knock down and which changes are secondary. To get around this problem we have adapted the auxin degron system for use in trypanosomes and here we describe its use in procyclic form *T. brucei*.

The auxin dependent degron is based on an Skp1-Cullin-F-box (SCF) E3 ubiquitin ligase that contains TIR1 as the F-box protein [7]. TIR1 determines specificity by binding to a substrate protein in an auxin (indole-3-acetic acid, IAA) dependent manner. The degron system was developed for use in yeast and mammalian cells and involved two steps. First, ectopic expression of TIR1 and second, the tagging of a target protein with an auxin induced degradation (AID) domain from the plant IAA17/AXR3-1 protein that is sufficient for binding to TIR1 in an auxin-dependent manner [7]. On the addition of exogenous auxin, the target protein is the ubiquitinylated and subsequently degraded by the proteosome (Figure 1). There is no known equivalent of the auxin dependent SCF in fungi or mammals and for the system to work the plant TIR1 has to bind the SKP1 component of the endogenous SCF. In addition, the endogenous levels of IAA in the experimental system have to be below the threshold for activating the SCF-TIR1.

**Figure 1.**
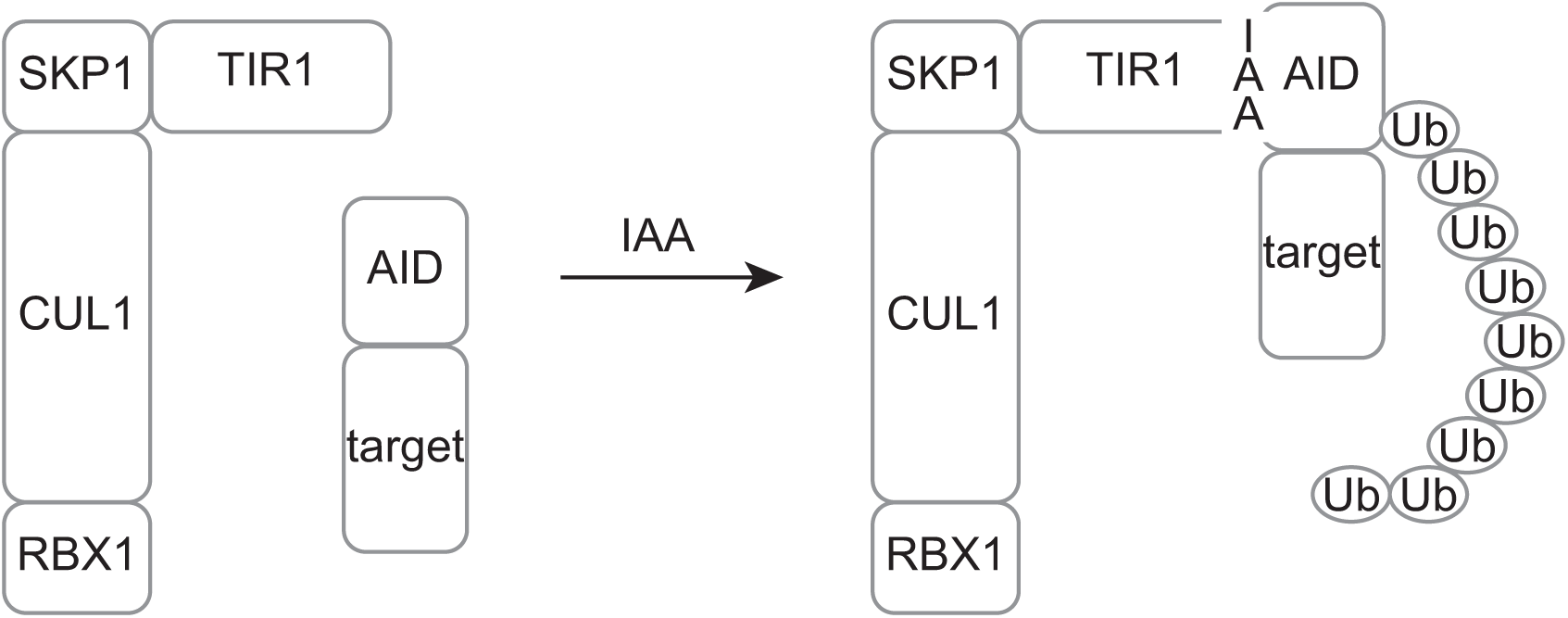
Cartoon illustrating the structure of the rice auxin-dependent E3 ubiquitin ligase and the binding of a protein tagged with an AID domain on the addition of the auxin indolyl acetic acid (IAA).

Initial experiments with procyclic form *T. brucei* in culture were unsuccessful, most likely due to a failure of TIR1 to interact with the endogenous SKP1. After a lot of faffing about, we decided to reconstruct the entire plant SCF-TIR1 in trypanosomes using the sequences of the components from rice as trypanosomes are cultured at 28°C or 37°C depending on the developmental stage. However, cell lines expressing these transgenes were only partially effective. Tagging of target proteins with AID domains in cell lines expressing the rice E3 ubiquitin ligase resulted in decreased expression levels in the absence of exogenous auxin. However, there was a glimmer of hope as addition of IAA resulted in degradation of the residual protein. The reduced level of the tagged protein is almost certainly due to levels of endogenous auxin produced by the metabolism of tryptophan [8]. Success followed the removal of this problem by altering the IAA binding site on TIR1 through changing phenylalanine 74 to glycine and using 5-phenyl IAA as the inducer [9].

The above serves as background to the methods below which describe first how to make a trypanosome cell line expressing rice SCF-TIR1 plus ARF, and second how to then produce a cell line with a specific gene tagged with a degron. The protocols are based on *T. brucei* Lister 427 procyclic forms but are applicable to other cell lines and developmental stages.

The open reading frames of four rice proteins, CUL1 (cullin/CDC53), SKP1, RBX1, and TIR1, were modified to include a single Ty epitope tag at the C-termini and were made synthetically after being codon optimised to increase expression in trypanosomes. In addition, a domain from rice auxin response factor 16 (ARF16) with an N-terminal Ty epitope tag was added to the mix as this had been reported to improve the functioning of the system [10]. The five transgenes were cloned into plasmid vectors in two sets: p4919 contains CUL1, SKP1, RBX1 and a G418 resistance marker; p5179 contains TIR1 F74->G, ARF16 fragment and a blasticidin resistance marker (Figure 2). Both plasmids contain PacI sites flanking the inserts and both were designed for read through RNA polymerase II transcription after integration into either the tubulin locus for p4919 or the PFR1 locus for p5179.

**Figure 2.**
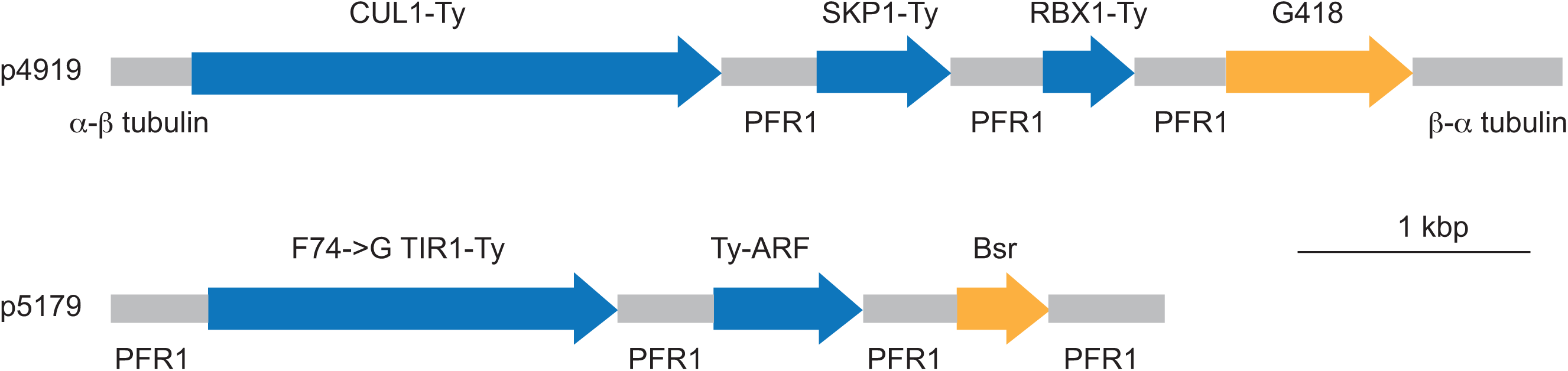
Cartoon showing the location of the genes encoding the E3 ubiquitin ligase components and a fragment of ARF16 in the plasmids p4919 and p5179. The fragemts show are released from the plasmids after PacI digestion. PFR1, paraflagellar rod protein 1.

To make a protein susceptible to auxin inducible degradation the gene is altered so that the encoded protein is a fusion with an AID domain. The method below is based on using derivatives of the pPOT plasmid series [11] as a template for making a PCR product that is used to modify the endogenous locus of the gene of interest (Figure 3). Trypanosomes are diploid and so this process is performed twice so both alleles are modified, one set of pPOT template plasmids encodes puromycin resistance, the second hygromycin. The success of the modification can be determined by western blotting either using an antibody recognising the protein of interest or epitope tags added as part of the modification of the endogenous gene and/or whole cell mass spectrometry.

**Figure 3.**
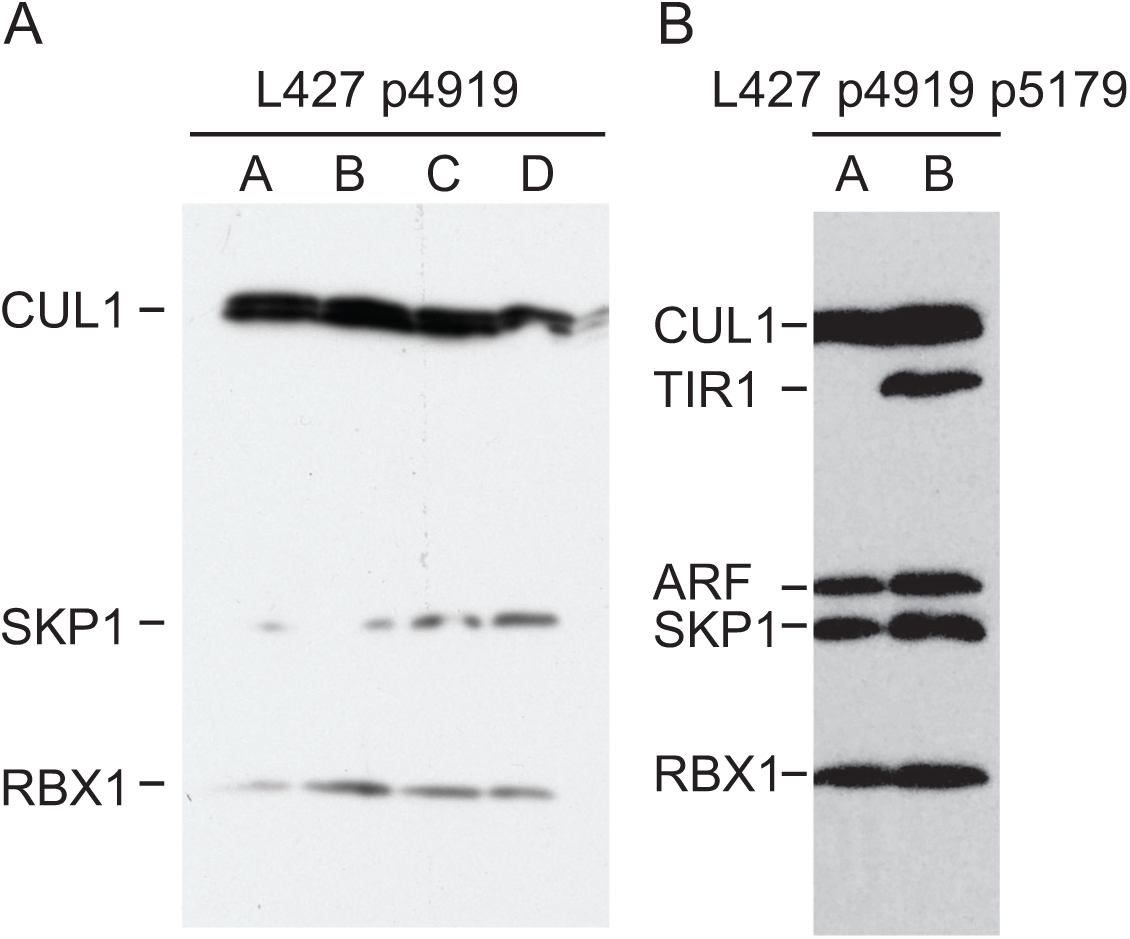
Western blots of screens for the expression of E3 ubiquitin ligase components. A. Four cell lines (A to D) isolated after transfection with p4919 and selection with G418. Whole cell lysates (1 x 10^6^ cell equivalents) were probed with anti-Ty and bands corresponding to the expected sizes of CUL1, SKP1 and RBX1 were observed. The strength of signal from CUL1 is always greater for unknown reasons and it is always a doublet, possibly due to modification with NEDD8. B. Two cell lines (A and B) isolated after transfection of cells with first p4919 and then p5179. Whole cell lysates (4 x 10^6^ cell equivalents) were probed with anti-Ty, cells in A did not express TIR1 and were not used further; cells in B successfully expressed all five proteins.

## 2 Materials

Follow local safety regulations when handling *Trypanosoma brucei*.

Prepare all solutions with ultrapure water and filter sterilize all solutions and store at 4oC.

### 2.1 Growth of procyclic form *T. brucei* in culture

1. *T. brucei* cell lines are available from www.atcc.org.
2. SDM79 medium plus 10% heat inactivated foetal bovine serum. SDM79 culture medium is available commercially.
3. Incubator for growth conditions: 27°C, humidified and 5% CO_2_.
4. Range of tissue culture flasks with vented caps.
5. 24 well plates for tissue culture.
6. Antibiotics for selection of transgenic cell lines 100x blasticidin: 1 mg/ml blasticidin 100x G418: 1.5 mg/ml G418 (geneticin) 100x hygromycin: 2.5 mg/ml hygromycin. 100x puromycin: 0.1 mg/ml puromycin.

### 2.2 Preparation of plasmids prior to electroporation

1. Plasmids p4919 and p5179 are available from Addgene (239357 and 239358)
2. PacI restriction enzyme and digestion buffer from New England Biolabs
3. Qiagen QIAquick PCR Purification Kit
4. 3 M sodium acetate buffer pH 5.0
5. Molecular biology grade ethanol
6. Material and equipment for agarose gel electrophoresis of DNA

### 2.3 Preparation of PCR products prior to electroporation

1. Template plasmids are available from Addgene: pPOTv7-puro-OsAID (239359), pPOTv7-hyg-OsAID (239360), pPOTv6-puro-3Ty:OsAID:3Ty (239361), pPOTv6-hyg-3Ty:OsAID:3Ty (239362), pPOTv7-hyg-AtAID:HA (239363).
2. 100-mer oligonucleotides
3. Standard PCR Thermocycler
4. Roche Expand PCR system
5. dNTP stock solution: 10 mM each dATP, dCTP, dGTP and dTTP
6. Material and equipment for agarose gel electrophoresis of DNA
7. Qiagen QIAquick PCR Purification Kit
8. 3 M sodium acetate buffer pH 5.0
9. Molecular biology grade ethanol

### 2.4 Electroporation to make transgenic cell lines

1. 15 ml and 50 ml sterile, conical and capped centrifuge tubes.
2. 20 ml syringes (sterile) and 0.22 μm syringe end filters (sterile)
3. Amaxa Nucleofector and 2 mm cuvettes.
4. Sterile molecular biology grade water
5. 3xR buffer: 200 mM Na_2_HPO_4_, 70 mM NaH_2_PO_4_, 15 mM KCl, 150 mM HEPES pH 7.3
6. Calcium chloride stock: 1.5 mM CaCl_2_
7. 50 mM 5-phenylindolylacetic acid in DMSO
8. Materials and equipment for western blotting
9. Monoclonal antibodies for detection of epitope tags: murine anti-Ty monoclonal antibody and rat anti-HA monoclonal, both commercially available

## 3 Methods

### 3.1 Growth of procyclic form *T. brucei*

1. Dilute the procyclic cells to 5 x 10^5^ /ml with SDM-79 medium and return to incubator.
2. Count the cell density after 24 and 48 hours. There should be a ∼5-fold increase in cell density per 24 hours.
3. After 48 hours dilute the cells to 5 x 10^5^ /ml and repeat.
4. Once the cells are proliferating at a constant rate, proceed to the electroporation. On the day before you plan to electroporate set up a 30 ml culture at 1 x 10^6^ /ml for each electroporation.

### 3.2 Preparing plasmids for electroporation

1. The day before the electroporation, prepare a PacI digest of p4919 or p5179. Digest 10 μg of plasmid with 50 U PacI in a 100 μl reaction following the manufacturers instruction. Incubate at 37°C for 2 hours and remove 3 μl for analysis by agarose gel electrophoresis, place the remainder of the digest at −20°C.
2. Analyse the digest using standard agarose gel electrophoresis for DNA. Both digests will contain two fragments: p4919: 2971 bp and 6236 bp; p5179: 2971 bp and 4501 bp.
3. The remaining 97 μl of the digested plasmid is then cleaned using a Qiagen QIAquick PCR Purification Kit and following the manufacturer’s instructions. Elute in 200 μl into a 1.5 ml tube, then add 20 μl 3 M sodium acetate and 550 μl ethanol, mix by shaking. Leave at −20°C overnight.

### 3.3 Electroporation and selection of cell lines expressing CUL1, SKP1 and RBX1

1. Recover the digested p4919 by centrifugation in a microfuge at 12000 rpm for 30 minutes, there should be a visible pellet.
2. It is now important to keep the digested plasmid sterile, so work in a sterile safety cabinet from here on.
3. Pour away the supernatant, close the tube and centrifuge again for 30 seconds, remove the residual supernatant with a 100 μl micropipette.
4. Dissolve the plasmid in 60 μl sterile water, add 35 μl 3xR buffer and 10 μl calcium chloride stock, mix by pipetting. Keep the dissolved plasmid in the sterile safety cabinet.
5. For each electroporation, prepare three T-75 culture flasks, flask 1 containing 26 ml fresh SDM79, flasks 2 and 3 containing 16 ml fresh SDM79.
6. Take the 30 ml procyclic form trypanosome culture and pipette 3 ml into a 15 ml centrifuge tube and 25 ml into a 50 ml centrifuge tube. Both tubes must have screw tops to keep the contents sterile.
7. Put the smaller tube aside for the moment.
8. Pellet the cells in the 50 ml tube at 3000 rpm for 5 minutes in a bench top centrifuge, return the tube to the laminar flow hood and recover just over 20 ml of the supernatant using a 20 ml syringe. Then add a 0.22 mm filter to the syringe and add 10 ml filtered conditioned medium to flasks 2 and 3.
9. Without delay, pellet the cells in the 15 ml centrifuge tube as above. Pour away the supernatant and centrifuge again for 30 seconds. Remove the residual supernatant with a 1 ml pipettor.
10. Resuspend the cells in 100 μl of the p4919 solution, transfer to a 2 mm cuvette, and electroporate using programme X-001. Immediately transfer the electroporated cells to flask 1 and swirl to disperse.
11. Transfer 1 ml from flask 1 to flask 2, swirl and then transfer 1 ml from flask 2 to flask 3. Place all three flasks in the growth incubator.
12. After 6 hours add 0.5 ml of 100x G418 stock to each flask (final 2x G418), swirl to mix and then pipette 1 ml aliquots into the wells of 24 well plates. The result is three 24 well plates with dilutions 1/1, 1/25 and 1/625.
13. Return the incubator and resist the temptation to examine the plates for 7 days.
14. After 7 to 10 days, selection will have occurred and it is likely that there will be some wells in the 1/625 dilution plate with proliferating cells and others with no cells. Inspect each well using an inverted microscope being careful to maintain sterility.
15. Mark wells with proliferating cells and note the number of wells.
16. If there are no wells with cells look at the 1/25 plate. All the wells in the 1/1 plate should contain proliferating cells, if not it is likely there were technical problems.
17. Once the cell density in a well is >5x10^6^ transfer the contents (of the well) into a T25 flask containing 10 ml fresh SDM79 with 0.1 ml G418 solution (1xG418). You should aim to pick wells from the highest dilution plate as these are more likely to be clonal. Pick four or more wells to analyse.
18. Passage the cells a couple more times (section 3.1) and move on to validation.

### 3.4 Validation

1. Prepare whole cell lysates for western blotting at 2x10^8^ cell equivalents/ml and analyse 10 or 20 μl.
2. Probe the western blot with anti-Ty monoclonal, typical results are shown in Figure 4A. Select a cell line in which the three Ty-tagged proteins of the expected molecular weights are expressed and proceed.

### 3.5 Electroporation and selection of cell lines expressing CUL1, SKP1, RBX1, TIR1 and ARF16 fragment

1. Proceed with the electroporation as above, start with a cell line successfully transfected with p4919 and grow in SDM79 supplemented with 1x G418 stock solution.
2. Use the PacI digested p5179 for the electroporation as in Section 3.3.
3. 6 hours after the electroporation, add 0.5 ml blasticidin stock (2x blasticidin) and 0.25 ml G418 stock per flask, then plate the three dilutions into 24 well plates as in section 3.3.
4. Be patient and once the cell density in a well is >5x10^6^, transfer the contents to a T25 flask containing 10 ml fresh SDM79 with 1x blasticidin and 1x G418. It is worth picking six or more wells to analyse and you should aim to pick wells from the highest dilution plate as these are more likely to be clonal.
5. Passage the cells a couple more times (section 3.1) and move on to validation (section 3.4) example results are shown in Figure 4B.
6. Select two cell lines to test for degron ability.

### 3.6 Synthesis of a PCR product for making degron cell lines

1. Design of oligonucleotide primers [11]. Two 100-mer primers are needed, the 5’ 80 nucleotides of each contains the sequence of the target site in the genome and, at the 3’ end, 20 nucleotides that base pair with the pPOT template. It should be noted that the quality of such long oligonucleotides varies with supplier.
2. PCR reaction. The choice of template plasmid will be determined by whether you would like to add epitope tags to your AID-tagged protein to determine depletion. You will need to assay for depletion and in the absence of specific assay or antibodies the use of an epitope tag works well.
3. We have used the Roche Expand system for these reactions, but other proof-reading thermostable polymerases should work as well.
4. Set up two reaction conditions for each amplification, one with 1.5 mM and the second with 2.5 mM MgCl_2_.

1.5 mM MgCl_2_:

1.5 mM $\text{MgCl}_2$ :
|  |  |
| --- | --- |
| pPOT template plasmid | 20 ng |
| Expand buffer 2 | 5 $\mu\text{l}$ |
| 10 mM each dNTP | 2 $\mu\text{l}$ |
| 10 pmol/microl oligo 1 | 1.5 $\mu\text{l}$ |
| 10 pmol/microl oligo 2 | 1.5 $\mu\text{l}$ |
| water | to 49 $\mu\text{l}$ |

2.5 mM MgCl_2_:

**Figure 4.**
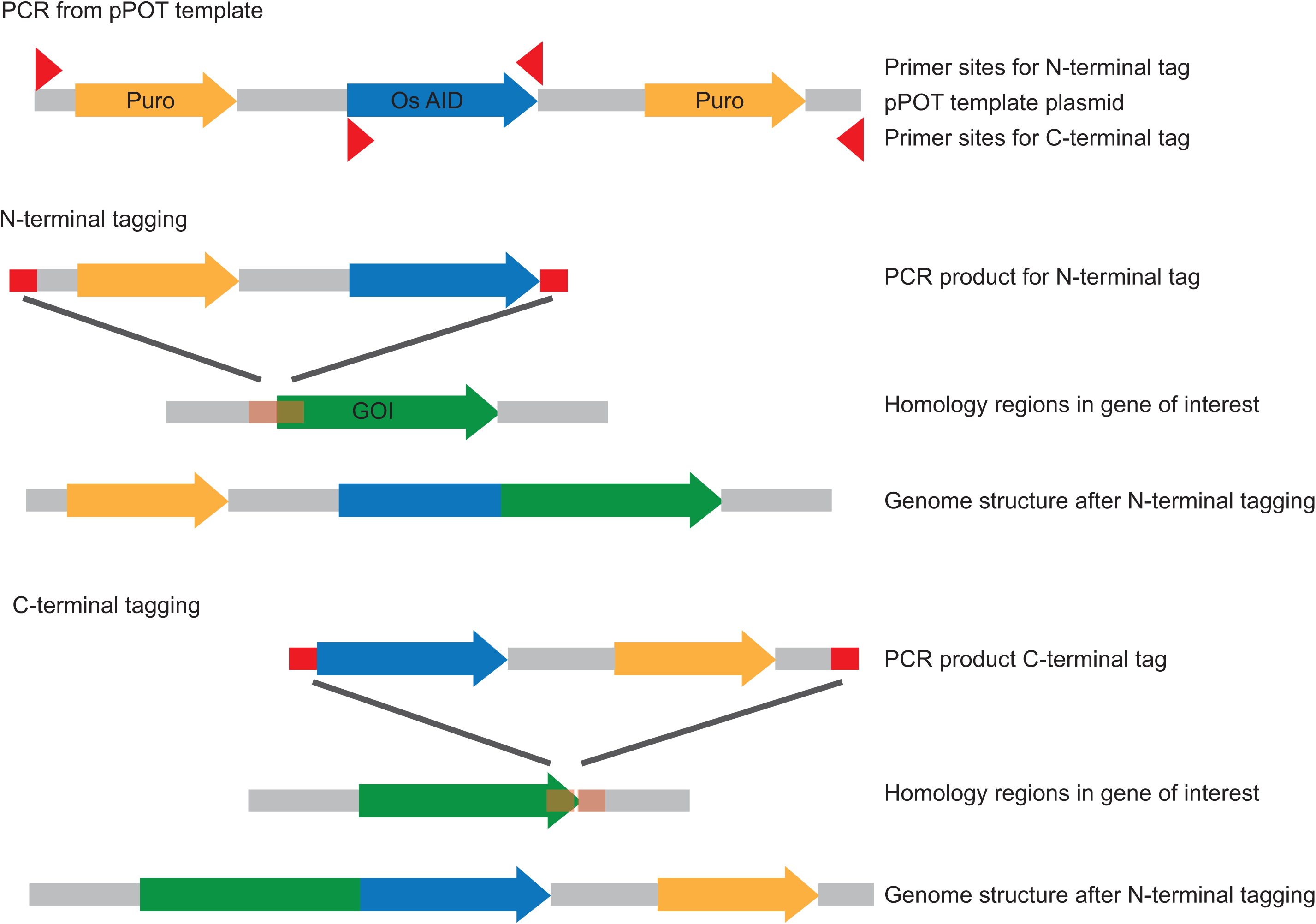
Cartoon showing principle behind use of pPOT plamids [11]. PCR products contain the tag, a selectable marker cassette plus 80 bp of targeting sequence at each end. The targeting sequences are in the 100mer primers used in the PCR.

5. Programme the thermocycler

6. Add 1 µl polymerase in the first cycle by pausing the programme once the temperature has decreased to 46°C.
7. At the end of the amplification reaction, analyse 2 µl of PCR product by agarose gel electrophoresis, store the remainder at −20°C.
8. You should expect a substantial amount of PCR product (Figure 5) and aim for 5 µg or more from the 50 µl reaction. Perform additional PCRs if necessary.
9. Purify the PCR product using a Qiagen QIAquick PCR Purification Kit. Elute in 200 ml into a 1.5 ml tube, then add 20 ml 3 M sodium acetate and 550 ml ethanol, mix by shaking. Leave at −20°C overnight.

2.5 mM $\text{MgCl}_2$ :
|  |  |
| --- | --- |
| pPOT template plasmid | 20 ng |
| Expand buffer 3 | 5 $\mu\text{l}$ |
| 25 mM $\text{MgCl}_2$ | 5 $\mu\text{l}$ |
| 10 mM each dNTP | 2 $\mu\text{l}$ |
| 10 pmol/microl oligo 1 | 1.5 $\mu\text{l}$ |
| 10 pmol/microl oligo 2 | 1.5 $\mu\text{l}$ |
| water | to 49 $\mu\text{l}$ |

5. Programme the thermocycler
|  |  |
| --- | --- |
| 96°C | 240 s |
| Cycles 1 to 5 |  |
| 94°C | 15 s |
| 46°C | 60 s |
| 68°C | 60 s per kbp of product |
| Cycles 6 to 30 |  |
| 94°C | 15 s |
| 60°C | 60 s |
| 68°C | 60 s + 5 s increase per cycle per kbp of product |
| After 30 cycles |  |
| 68°C | 420 s |
| 4°C | indefinite |

### 3.7 Electroporation to tag the first allele

1. Culture the degron proficient cell line in SDM-79 with 1x G418 and 1x blasticidin, once it is proliferating at the expected rate, prepare 30 ml cultures at 1x10^6^ /ml the day before the electroporation.
2. Recover the PCR product by centrifugation in a microfuge at 12000 rpm for 30 minutes as in Section 3.2. There should be a visible pellet.
3. Dissolve as for the plasmid in Section 3.2.
4. For each electroporation, prepare three T-75 culture flasks, flask 1 containing 28 ml fresh SDM79, flasks 2 and 3 containing 16 ml fresh SDM79.
5. Proceed as in section 3.3 and add 10 ml conditioned medium to flasks 2 and 3.
6. Perform the electroporation as in section 3.3 and add the cells to flask 1. Transfer 2.5 ml from flask 1 to flask 2, swirl to mix then transfer 2.5 ml from flask 2 to flask 3 so you end up with 1 in 10 and 1 in 100 dilutions.
7. After 6 hours, either add 0.5 ml puromycin stock (2x puromycin) or 0.5 ml hygromycin stock (2x hygromycin) to flasks 1, 2 and 3 and plate out in 24 well plates. The choice of puromycin or hygromycin is determined by the template plasmid used to make the PCR product.
8. Return to the incubator and after 7 to 10 days check each well for active proliferating cells, you should expect between 20 and 500 transformants per electroporation.
9. Once the cell density in a well is >5x10^6^, transfer the contents to a T25 flask containing 10 ml fresh SDM79 with 1x blasticidin, 1x G418 and 1x puromycin/hygromycin. It is worth picking six or more wells to analyse and you should aim to pick wells from the highest dilution plate as these are more likely to be clonal.
10. Passage the cells a couple more times (section 3.1) and move on to validation (section 3.4) example results are shown in Figure XX.

### 3.8 Validation of degron cell lines

1. Prepare whole cell lysates for western blotting at 2x10^8^ cell equivalents/ml and analyse 10 or 20 μl. Make lysates before and 1 hour after the addition of 5Phe-IAA to 50 µM.
2. Probe the western blot with either a protein specific antibody or anti-Ty /anti-HA monoclonal if you have added an epitope tag.
3. With a protein specific antibody, you should observe a band corresponding to the untagged protein and one for the tagged protein. The tagged protein band should be diminished on addition of the inducer (Figure 6A).
4. In the case of using a monoclonal, you should see a single band that goes after addition of inducer. The use of a Ty-AID for the tagging of the first allele has an advantage as it confirms the E3 ubiquitin ligase components are still being expressed (Figure 6B).
5. Select a couple of cell lines in which the tagged protein is degraded after addition of inducer.

### 3.9 Electroporation of tag the second allele

1. Grow the cells from with antibiotic selection, 1x G418, 1x blasticidin and 1x puromycin or hygromycin.
2. Repeat the process described in Section 3.7 and select 4 to 6 wells from the most dilute plate. These should now proliferate happily in the presence of all four antibiotics.

### 3.10 Validation of degron cell lines

1. Again, grow the cell lines with all four selection antibiotics at and check for degradation of the target protein after addition of inducer by western blotting as in section 3.8.
2. If a protein specific antibody is available, then the western blot will show if both alleles are tagged and whether the protein is degraded on addition of inducer.
3. In the absence of a specific antibody, probe the blots sequentially with anti-HA, then anti-Ty, and then a third antibody as a loading control (Figure 6B). This provides evidence that both alleles are tagged and degraded.

### 3.11 Validation by mass spectroscopy

A second approach is to use whole cell mass spectrometry before and after addition of 5Phe-IAA although this may not be sensitive enough with low copy number proteins.

**Figure 5.**
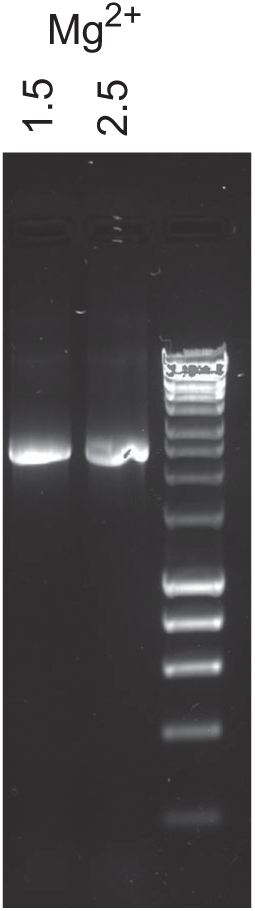
Example of the PCR products used to add the AID tag by homologous recombination at the endogenous allele. The products made with 1.5 and 2.5 mM Mg2+ are indicated and 2 μl of the PCR reaction were analysed.

**Figure 6.**
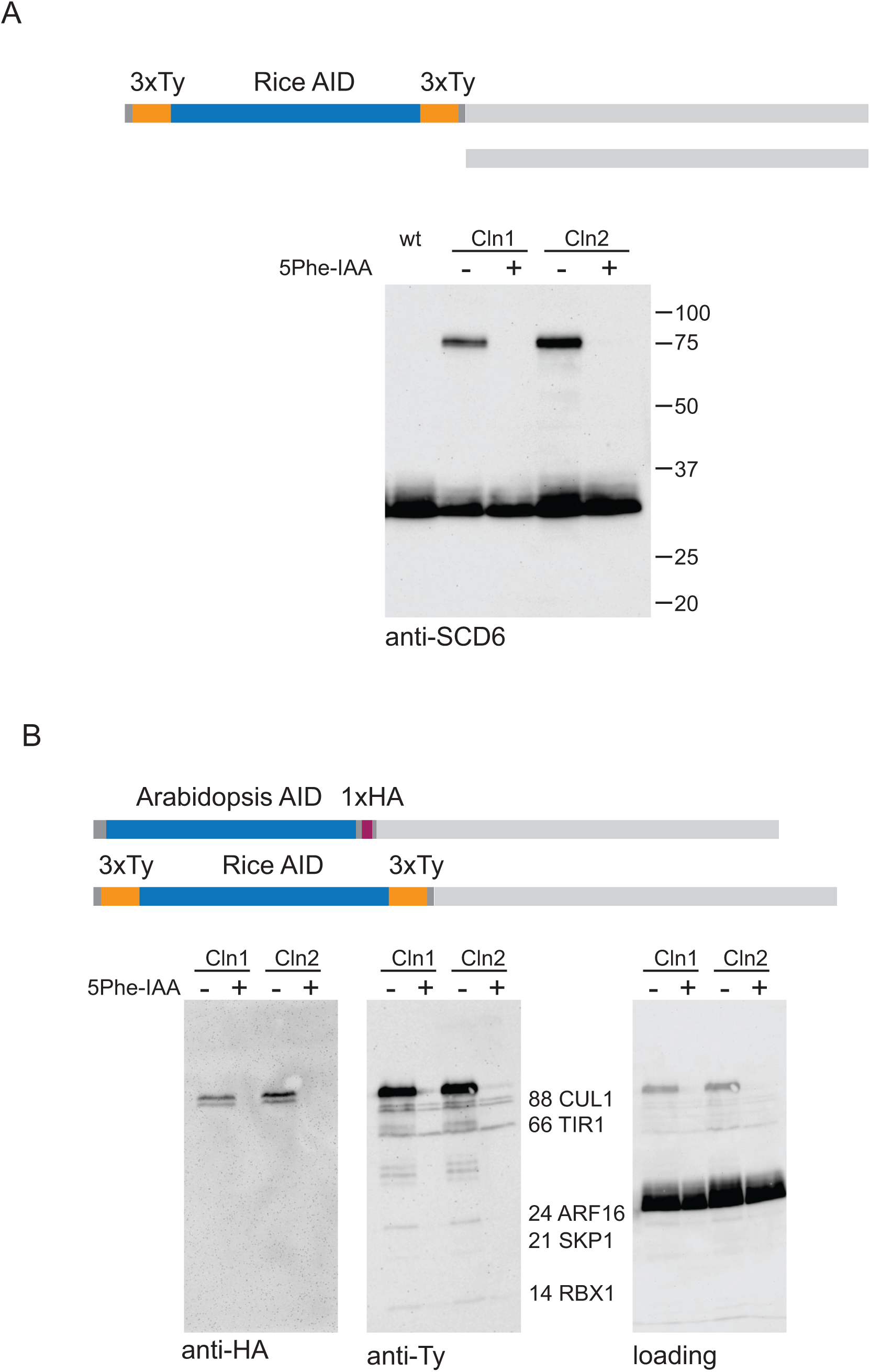
A. Cartoon showing the two alleles of the target gene and a western blot probed with anti-SCD6 showing successful tagging of one allele of the SCD6 gene and degradation of the tagged protein 1 hour after the addition of 50 μM 5PheIAA. 1x10^6^ cell equivalents loaded per track. B. Cartoon showing the two alleles of target gene and western blots illustrating the use of epitope tags to demonstrate tagging of both alleles and degradation of the targeted protein. 2 x 10^6^ cell equivalents were loaded per track. One allele was modified with an Arabidopsis AID domain containing a single HA epitope, the second was modified with a Rice AID domain containing six Ty epitopes. Both are degraded after addition of 5PheIAA.

### 3.12 You are now in a position to do the interesting experiments

## 4 Notes

1. The rice E3 ubiquitin ligase is present in the cytoplasm and so will only work with those accessible proteins, probably just the cytoplasm and nucleus. It will not work for proteins inside the endosome. At this stage, success with proteins in transit to the mitochondrion or transmembrane proteins with a small cytoplasmic domain remains unknown. We have used the degron cell line with around 15 different cytoplasmic proteins and all have been successfully degraded on induction.
2. The approach is dependent on the AID tag being tolerated by the cell.
3. The system should be easily transferable to other trypanosomatid species.

## Acknowledgements

MC is a Wellcome Investigator (217138/Z/19/Z). RPdL CR and DNM were funded by a CNPq Post-doctoral international fellowship in the Science Without Borders Program. JdFN was funded by CAPES and Cambridge Overseas Trust. AM was funded be a Wellcome summer studentship. AK was funded by St John’s College.

